# Cell Fate by Design or Chance: Interplay of *Nonlinearity* and *Stochasticity* in Gene Regulatory Network motifs

**DOI:** 10.1101/2025.09.17.676733

**Authors:** Mrinal Sarkar, Ushasi Roy

## Abstract

Cell fate decisions emerge from the intricate tug-of-war between deterministic regulatory logic and stochastic molecular fluctuations. How stochasticity, inherent in intracellular Gene Regulatory Networks (the heart of determining cellular fates), shapes cellular regulatory dynamics is central to and essential for both interpreting natural cellular decision-making and designing robust synthetic circuits. In this work, we develop a general mathematical framework based on the linear noise approximation (LNA) to quantify intrinsic fluctuations around the mean expression levels in a single-cell GRN motif with nonlinear cooperative regulatory couplings. The genetic toggle switch (TS) and TS coupled with positive autoregulation, as illustrative examples, exhibit both continuous and discontinuous transitions (bifurcation) between low- and high-expression states driven by nonlinear regulation. We demonstrate that LNA captures both deterministic and stochastic dynamics. We find that the LNA performs well in capturing the inherent stochasticity in both monostable and bistable regimes. Still, deviations arise near bifurcation points: fluctuations are enhanced near criticality in continuous transitions, while finite-size effects dominate discontinuous transitions, analogous to critical phenomena in statistical physics. Finally, we discuss the implications of our framework for designing synthetic gene circuits.

## I. INTRODUCTION

Gene Regulatory Networks (GRNs), laying out the blueprint for gene expression, along with multiple regulatory mechanisms, epigenetic alterations, signaling cascading dynamics, and environmental cues, govern the identity, functions, and behavior of a biological cell. GRNs consist of interconnected genes and their products, including transcription factors (TFs), RNA molecules, and proteins, that regulate one another’s expression through various molecular interactions. GRNs determine a wide range of biological phenomena, from cell differentiation, development, and tissue homeostasis to stress responses, metabolic regulation, and disease progression.

Despite the complex architecture of hundreds and thousands of interacting components, GRNs, fundamental to orchestrating gene expression, often saliently feature simple 2-, 3-node recurring motifs manifesting emergent multistable and oscillatory dynamics, robust to intrinsic and extrinsic noise, playing a crucial role in various cell-fate decisions [1, 2]. Multistability, a key feature central to differentiation, adaptation, and phenotypic heterogeneity, enables cells with identical genotypes to have different stable phenotypes [3]. Inherently stochastic bio-molecular processes can perturb these deterministic stable phenotypic states, influencing the final cellular fates (stable states) and switching between these fates.

Nonlinearities and inherent stochasticities in gene expression (both transcription and translation) are two fundamental aspects of how genes are regulated and expressed in biological systems. Both play key roles in determining the behavior of gene regulatory networks (GRNs), and their interplay can produce rich and diverse cellular phenotypes. Inherent nonlinearities at the intracellular genetic level drive the emergence of multistable and oscillatory behavior at the cellular level. Nonlinearities at the gene level refer to the complex, non-additive relationships between genes and their regulatory elements, such as transcription factors, repressors, and enhancers. The reason for underlying nonlinearities can be manifold – for instance, feedback loops, cooperativity, nonlinear synthesis and degradation rates, etc. At the cellular level, these gene-level nonlinearities create multiple stable gene expression patterns, or attractors, in the system’s phase space.

Stochasticity in gene expression [4–7], on the other hand, stems from both intrinsic sources—such as the random timing of transcription and translation events and the low copy number of molecules—and extrinsic sources like fluctuations in cellular components (e.g., RNA polymerases, ribosomes) and environmental variability. These random fluctuations lead to significant variability in gene expression levels across cells, even within a genetically identical population under uniform conditions. This noise plays a critical role in enabling phenotypic heterogeneity, driving cell fate decisions during development, and supporting bet-hedging strategies that a population often undertakes to increase its resilience to environmental stress.

One way of quantifying the inherent stochasticity in these processes is by calculating the Fano Factor (FF). It is defined as the variance-to-mean ratio in the copy number of the relevant molecular species. An FF value of unity indicates a Poisson process. FF greater than unity indicates over-dispersion relative to Poisson noise, reflecting enhanced stochasticity that may be biologically significant in modulating multistable dynamics and therefore cell fate decisions.

From a synthetic biology perspective, most gene circuits are designed and analyzed using deterministic descriptions, typically formulated as systems of coupled ordinary differential equations (ODEs) [8, 9]. Such models provide a ***mesoscopic*** *description*, in which the dynamics is expressed in terms of *average molecular concentrations*, rather than the discrete copy numbers of genes or proteins considered in a ***microscopic*** *description*. While these ODE models capture nonlinear interactions and yield fixed point solutions, they neglect intrinsic stochasticity. As a result, they exhibit bifurcation behaviors, attractor solutions, and multistability, but fail to address the *fluctuations* around deterministic solutions. In practice, however, fluctuations around each fixed point can be crucial, especially in regimes of multistability, where noise may determine whether a designed state is stably maintained or whether switching between states is induced. Quantifying these fluctuations around attractors, for instance through the *Fano factor*, provides a natural measure of circuit reliability [6, 10]. Furthermore, understanding noise characteristics is essential for the rational design of synthetic oscillators, toggle switches, and logic gates, since their performance depends not only on mean expression levels but also on the full distribution of molecular states [5, 11]. A systematic characterization of fluctuations is therefore necessary to bridge deterministic circuit design with the inherently stochastic nature of gene expression, which constitutes the central motivation of the present work.

Gene regulatory networks (GRNs), in the *macroscopic description*, are commonly modeled using production and degradation rates to describe the time evolution of protein concentrations. The *production rate* of a protein incorporates both the *transcription rate* of the corresponding gene (mRNA synthesis) and the *translation rate* of mRNA into protein, while the degradation rate captures *protein decay* due to dilution or enzymatic degradation [10, 12]. In this framework, gene *A* (protein concentration [*A*]) evolves according to *d*[*A*]*/dt* = *g*_*A*_ −*γ*_*A*_[*A*], where *g*_*A*_ represents the *effective* production rate and *γ*_*A*_ the degradation rate. *Transcriptional regulation* by *other genes* is often incorporated via **nonlinear functions**, such as Hill functions, which modulate *g*_*A*_ according to the concentrations of regulators. For instance, for a gene *A* regulated by a gene *B*, the dynamics of gene *A* now follows *d*[*A*]*/dt* = *g*_*A*_*H*(*B*) −*γ*_*A*_[*A*], where *H*(*B*) is a Hill function representing the regulatory effect of *B* on *A*, with specific details provided in the main text. At the microscopic level, the continuous ‘concentration’ variable is replaced by discrete molecule numbers. Production and degradation events occur probabilistically, with propensities determined by the corresponding rates. The stochastic dynamics generate trajectories of molecule counts over time, capturing intrinsic noise arising from the discrete and random nature of biochemical reactions.

In this work, we consider a general *M* -node gene regulatory network (GRN) inside a cell, where each gene interacts through *nonlinear Hill functions* and possesses its own degradation rate. We analyze both the macroscopic and microscopic descriptions of the system. Starting from the chemical master equation (microscopic perspective), we develop a general framework and apply the standard *Linear Noise Approximation (LNA)* to derive the *mesoscopic dynamics*, which allows us to quantify fluctuations (variances and covariances) around the mean expression levels (deterministic fixed points). A major finding is the crucial role of inherent stochasticity in cooperative gene behavior and its quantification through the Fano factor. We show that deterministic dynamics are sufficient to estimate the variance of protein levels, providing an explicit link between mean behavior and noise. We show explicitly that it is the inherent stochasticity manifested through the nonlinear Hill function that drives the Fano factor away from unity, leading to non-Poissonian behavior. Using a simple motif (the toggle switch), we verify our analytical predictions: they agree well in the monostable regime, but deviations appear in the critical regime (near bifurcation points) of (bi)multistability, highlighting the need for caution near bifurcation points, particularly when designing synthetic circuits.

The LNA provides a systematic framework to quantify stochastic fluctuations around the deterministic mean of a biochemical system [13, 14]. By expanding the chemical master equation around the macroscopic steady state, the LNA yields a Gaussian approximation for the probability distribution of molecular concentrations, enabling computation of variances and covariances without resorting to full stochastic simulations [10, 15]. This approach is particularly useful for GRNs, where molecule numbers are moderate and intrinsic noise plays a significant role, but exact solutions of the master equation are intractable. While previous studies using the LNA have mainly focused on **linearly coupled gene expression models** [10, 14, 15], here we explicitly apply it to **general** *M* **-node GRNs with nonlinear Hill-function interactions**. This enables us to uncover how **intrinsic stochasticity interacts with nonlinear cooperative regulation**, a question that has not been systematically addressed in prior work. Our framework is very general in that it applies to any *M* -node GRN, thereby extending previous LNA analyses from linear or small networks to fully nonlinear GRNs. These results provide a foundation for understanding noise in natural systems and for designing synthetic genetic circuits with predictable fluctuation characteristics.

Note that by *macroscopic limit* we mean the infinite system-size limit, where fluctuations become negligible and the GRN dynamics reduces to deterministic rate equations for gene concentrations [16]. On the contrary, at the *mesoscopic level*, fluctuations are small but non-negligible, accounting for intrinsic noise around these *macroscopic concentrations*, and correspond to a finite system-size description.

The paper is organized as follows. Sec. II describes the model and provides an overview of the GRN network. In Sec. III, we develop the general framework for an *M* -node GRN motif using the Linear Noise Approximation (LNA).

Sec. IV applies this framework to specific cases, namely, the Toggle Switch (TS) and Toggle Switch with self-activation (TSSA), with explicit computation of FFs. In Sec. V, we validate our theory through deterministic and stochastic Gillespie simulation results along with a discussion of the findings. Sec. VI finally concludes with a summary and outlook. Detailed derivation of the FFs for the TS is provided in Appendix A.

## II. MODEL DESCRIPTION

We illustrate the fundamental features of our model via the schematic (see Fig.1). A *complex network* of cells interacting with one another is depicted on the Left. The nodes (blue and red filled circles) represent the cells, and the edges represent the interactions (intercellular signaling events). The specific *red* -filled cell with the intracellular genetic interactions (in the form of TS for illustration; it can be any other motifs) is shown at the Center. The discrete dynamics of one of the genes (of type A) is shown at the right. Interactions at the intercellular level (e.g., signaling among cells) can be considered by establishing links from one node to another, and thus, we construct the network. Each cell can have its own GRN motif, and hence the intracellular discrete dynamics. The model we develop here is very general and phenomenological, since it has the strength to capture all the key attributes of a many-component interacting system, with different types of entities influencing positively or negatively one another’s downstream regulatory pathways, with underlying nonlinearities and stochasticity. The *complex network* of cells with intracellular genetic regulation in terms of motifs may be completely or partially connected.

In this study, we consider the *intracellular genetic regulatory dynamics* of an isolated cell. The rightmost part of the figure illustrates the stochastic dynamics of gene-A involving discrete random variables *N*_*A*_, which represents molecule number of gene-A. At any time *t*, the system is described by the probability distribution *P* (…, *N*_*A*_, …; *t*), where *N*_*A*_ denotes the copy number of species A. The dynamics consists of two opposing stochastic transitions: *N*_*A*_ → *N*_*A*_ ±1, where the +1 transition corresponds to *production* with rate *g*_*A*_ℋ (*ϕ*_*B*_), and the −1 transition corresponds to degradation with rate *γ*_*A*_ (proportional to *N*_*A*_). The production rate depends on a regulatory input *ϕ*_*B*_∝ *N*_*B*_ passed through a nonlinear Hill function ℋ, reflecting how species B influences species A, through gene regulation. We develop the mathematical formalism for studying the stochastic dynamics of a GRN motif embedded inside an isolated cell. Every cell has the same stochastic dynamics. Note that the cell-level network (nodes: cells, links: cell-to-cell signaling), as shown in Fig. 1, is included to only illustrate the scenario; the present analysis focuses on isolated intracellular dynamics, while the former (cell-level network) will be the subject of future work.

**FIG. 1.**
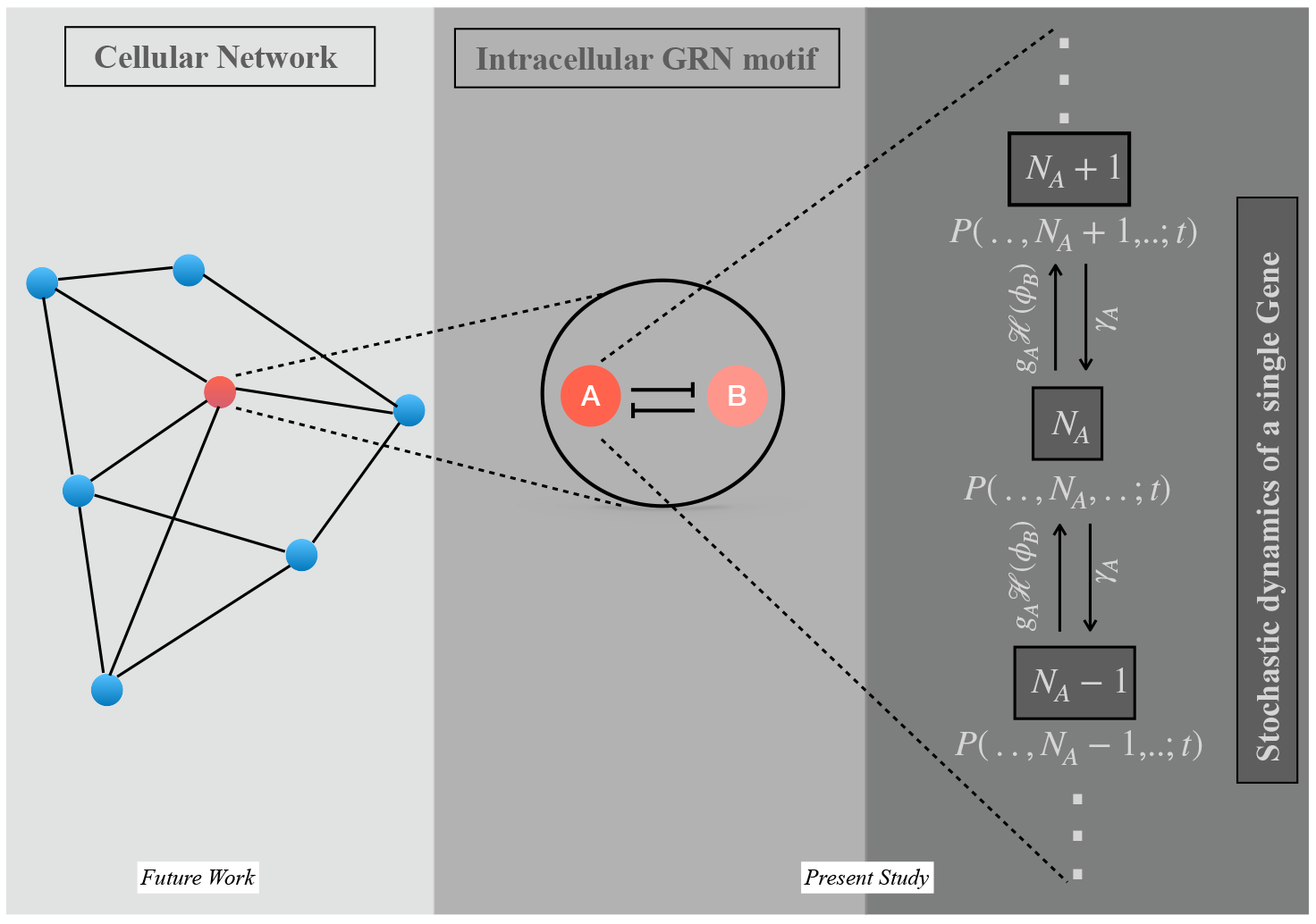
Schematic representation of the model. **Left**: A *complex network* of intersacting cells. (This is shown for the illustration of the whole scenario. (The detailed analyses would be executed in a future study!) **Center**: A gene regulatory network (GRN) motif (depicted by a toggle switch (TS) for illustration) within an isolated cell (red filled circle). **Right**: A zoomed-in view of the discrete stochastic dynamics of a certain type of gene A, interacting with gene B, in the TS within the isolated cell. It illustrates how the copy number of A-type genes, *N*_*A*_, evolves stochastically in time due to the interplay of *production* and *degradation* events. The gray boxes indicate the copy numbers (*N*_*A*_, *N*_*A*_*±* 1), and the arrows denote the possible transitions with their respective rates. Each cell in the network exhibits similar intrinsic stochastic dynamics.

## III. MATHEMATICAL FRAMEWORK

This section develops a general formalism for the Linear Noise Approximation (LNA) of a GRN motif with *M* genes, embedded inside a biological cell, following [13]. To this end, consider an *M* -node GRN motif, where each type of gene, say, the *i*-th one, affects (activates or inhibits) the other *j*-type of directly connected genes, where *j* ≠ *i*; *j* = 1, 2, 3, · · ·, *M*. From now on, we will use the term *i*-genes to denote the *i*-th type of genes for convenience.

### A. Macroscopic dynamics

Let *ϕ*_*i*_ be the concentration (a macroscopic quantity, describing the behavior) of the *i*-th gene in a *M* -node GRN, and define ***ϕ*** = (*ϕ*_1_, *ϕ*_2_, *ϕ*_3_, · · ·, *ϕ*_*M*_)^**T**^. The deterministic dynamics of the *i*-th gene are governed by

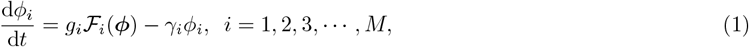

where ***ℱ*** = (ℱ_1_, ℱ_2_, ℱ_3_, · · ·, ℱ_*M*_)^**T**^ and 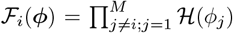. Clearly, ℱ_*i*_(***ϕ***)s are nonlinear functions in ***ϕ***. The parameters *g*_*i*_ and *γ*_*i*_ are respectively the production and degradation rates of *i*-th gene, and the product term represents the interaction with *all other genes*, with ℋ (*ϕ*_*j*_) being the shifted Hill function corresponding to the *j*-th gene. Note that we consider the *all-to-all* case, where each gene in the network influences every other gene. A more general scenario could be described by introducing an adjacency matrix in ***ℱ***, with elements *a*_*ij*_ = 1 if genes of types *i* and *j* are connected and *a*_*ij*_ = 0 otherwise. However, this generalization is beyond the scope of the current study.

The Hill function ℋ(*ϕ*_*j*_) is defined as follows:

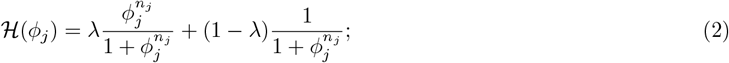

where *n*_*j*_ represents the Hill coefficient describing the cooperativity of other *j* genes, referring to how the binding of one molecule affects the likelihood of additional molecules binding to the same gene regulatory region of *i*-gene. The parameter *λ* is the fold change in the levels of the associated proteins due to transcriptional regulation: *λ* = 0 refers to repression/inhibition while *λ* = 1 denotes activation. The number of shifted Hill functions in the first term in the RHS of Eq.1 depends on the number of incoming links to the *i*-th gene, i.e., the number of other genes regulating the expression of the *i*-th gene. The Hill function is widely used in modeling gene regulatory networks because it provides a simple yet powerful mathematical function to capture nonlinear, cooperative binding behavior between transcription factors and DNA. To keep the model generic, we employ the Shifted Hill Function ℋ(*ϕ*_*j*_), a general, widely-used mathematical form to capture transcriptional cooperative regulation by activators or repressors depending on the signs of fold change *λ* [17–19].

### B. Microscopic dynamics

We now switch from the macroscopic continuum concentration to the microscopic discrete number basis to understand the system in greater detail. Define *P* (***N***, *t*), where ***N*** = (*N*_1_, *N*_2_, *N*_3_, · · ·, *N*_*M*_)^**T**^, to be the conditional joint probability of having *N*_*i*_ type-*i* genes (*N*_1_ type-1 genes, *N*_2_ type-2 genes, and so on) at any given instant of time *t*, given that there were *N*_*i*_ = *N*_*i*0_ at the initial time *t* = 0, where *i* = 1, 2, 3, · · ·, *M* are the different types of interacting genes. If Ω is the total volume of the cell, one can write down the following master equation [13]:

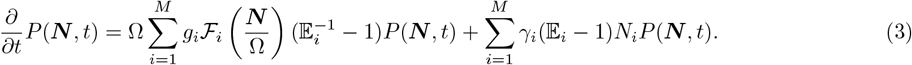

Here 𝔼_*i*_ is the step operator that acts in the number/discrete basis as [13]:

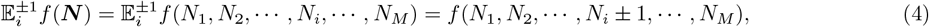

for any function *f* which depends on the number of different types of genes, and this number can get incremented/decremented depending on being operated by the step operator 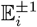. We would now proceed with a systematic expansion in Ω^−1*/*2^ with an expectation that, in its lowest order, this method would reproduce the deterministic macroscopic dynamics.

*Ansatz for system-size expansion*: By definition, the macroscopic concentration of *i*-gene *ϕ*_*i*_ = lim_Ω*→∞*_(*N*_*i*_*/*Ω). At the mesoscopic level, assume *ξ*_*i*_ be the fluctuation around *ϕ*_*i*_, we take the ansatz 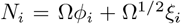. In vector notation,

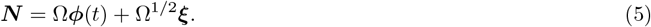

The joint probability can now be expressed in terms of new ‘intensive’ variables

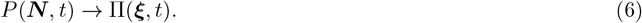

As it is clear from the above equation, the probability *P* is a function of the number of genes, while Π is a function of the intrinsic fluctuations in the number of genes involved.

Equation (6), on taking the time derivative at a constant ***N***, and using Eq. (5) yields,

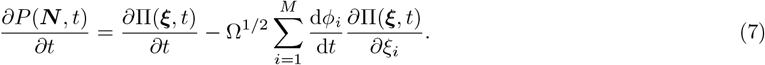

The step operator in Eq. (4) acts in the concentration basis (**mesoscopic** scale) as a change from *ξ*_*i*_ to 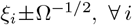. The fact that

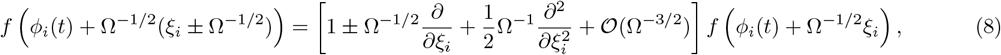

implies, for each *i*,

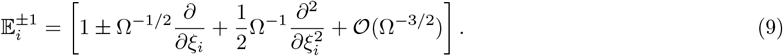

Now, the transition rate function ℱ_*i*_ takes the form under system-size expansion (by expanding in a Taylor series around the macroscopic mean *i.e*. concentration ***ϕ***) :

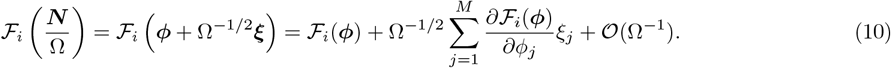

Note that the rate function ℱ_*i*_(***ϕ***) involving the product of nonlinear Hill functions in the macroscopic equation) does not include *ϕ*_*i*_, and hence the second term in the above Eq. (10) automatically respects the condition, i.e., *j*≠*i*.

Substituting Eqs. (5), (7), (9) and (10) in Eq. (3), we obtain

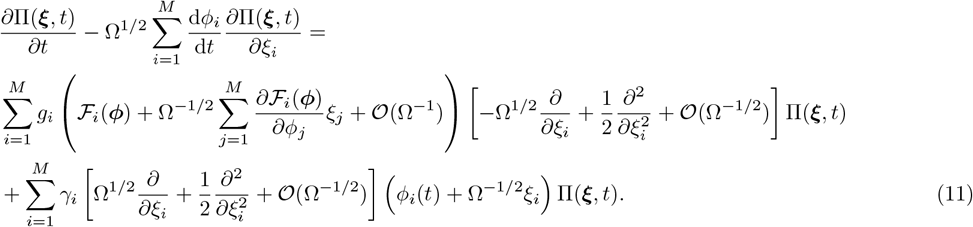

We have to match coefficients of successive powers of Ω from both sides of Eq. (11). Equating the coefficients of Ω^1*/*2^ we obtain

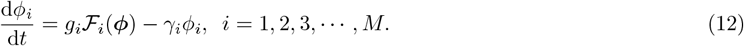

We thus recover the macroscopic equation, same as Equation (1) for ***ϕ***, as expected. Similarly, equating the coefficients of Ω^0^ yields

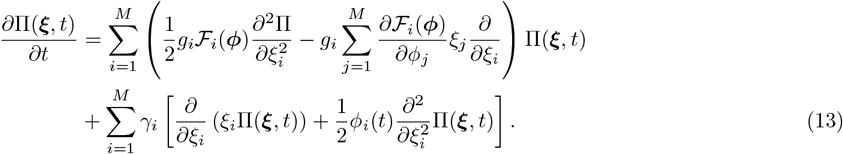

On rearranging, this yields,

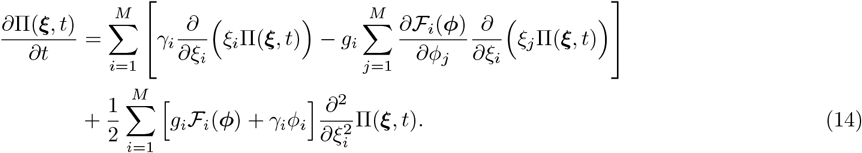

Equation (14) can be cast in the following form:

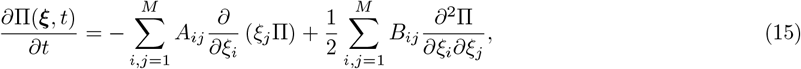

with initial conditions 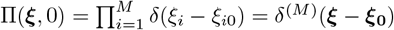. Here

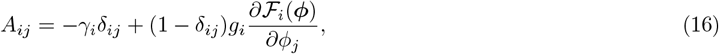

and ***B*** is a diagonal matrix with elements given by

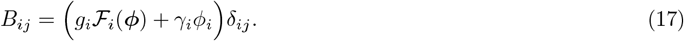

Equation (15) is a linear multivariate Fokker-Planck equation (FPE) for Π(***ξ***, *t*) whose solution is a Gaussian distribution given by [13, 20]

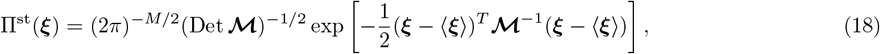

where 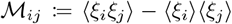. The above solution, given by Eq. (18), being Gaussian, all higher moments of the variables, except for the first two, vanish. We are interested in the behavior of the non-vanishing moments only, namely, the mean and variance, which can be computed from Eq. (15). One can show that the stationary-state mean and the covariance matrix ℳ satisfy the Lyapunov equation [13]

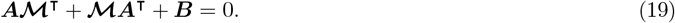

### C. Fluctuations: Mean and variance

One can thus compute auto- and cross-correlations in the intrinsic number fluctuation around the macroscopic concentration of the *M* different types of genes from Eq. (15). Multiplying Eq. (15) by *ξ*_*k*_ and integrating over all *ξ*, one obtains the equation for the temporal evolution of the first moment (mean)

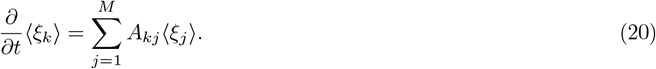

Similarly, the equation for the temporal evolution of the second moment (variance) is obtained by multiplying Eq. (15) by *ξ*_*k*_*ξ*_*l*_ and integrating over all *ξ*,

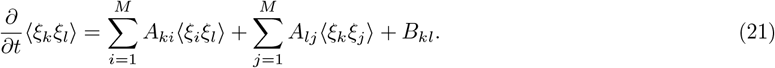

This gives us the auto- and cross-correlations in the intrinsic number fluctuation of the total number of different types of *M* -genes. In the next section, we apply these LNAs explicitly to specific cases of two genes and investigate the role of intrinsic fluctuations.

### D. An equivalent stochastic differential equation (OU process)

Before proceeding to study specific cases, it is worth mentioning that the Fokker-Planck equation for the fluctuation, Eq. (15), can be equivalently expressed as an Itô stochastic differential equation of Ornstein–Uhlenbeck type:

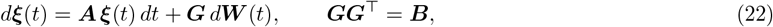

where ***W*** (*t*) is an *M* -dimensional Wiener process. Since ***B*** is diagonal and positive definite [Eq. (17)], one convenient choice is

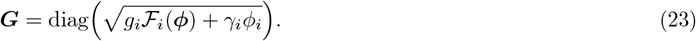

Here ***G*** matrix elements represent the amplitude of the noise in the system. In the OU process Eq.(22):

1. The drift term ***Aξ*** corresponds to the linearized deterministic dynamics around the macroscopic steady state.
2. The diffusive term ***G*** *d****W*** (*t*) captures the stochastic fluctuations, with covariance *B* = *GG*^*T*^.

The intrinsic fluctuations can thus be interpreted as a deterministic dynamical system subjected to Gaussian noise, valid in the vicinity of the macroscopic steady state. Equivalently, the LNA maps the intrinsic fluctuations of a general *M* -node GRN onto a multivariate OU process, with Gaussian fluctuations fully characterized by (*A, B*).

## IV. APPLICATIONS OF OUR GENERAL MATHEMATICAL FRAMEWORK

We will commence this section with the illustrative example of a classic and foundational two-component motif in the study of gene regulatory networks (GRNs), viz., the Toggle Switch (TS). Then we will add more components and interactions to the simple network architecture, to study the implications of nonlinearities and stochasticities in each specific case, and the differences that it incurs from the basic case of TS.

### A. Motif 1: Toggle Switch

In this section, we take up the specific case of a two-component GRN motif, viz., Toggle Switch (TS). This is a natural choice of the simplest multi-component motif, capable of exhibiting bistability (multistability at its lowest rank). A TS consists of two genes (or gene products) that mutually repress each other. This simple architecture can give rise to bistability, where the system can settle into one of the two stable expression states (High A /Low B or Low A /High B). Mutual repression creates a feedback loop where, if A is highly expressed, it suppresses B, reinforcing its dominance, and high expression of B suppresses A, strengthening its expression. This is one of the simplest motifs for bistability, exhibiting hysteresis, cellular memory, noise, and switching. It captures binary cell fate decisions — such as bacterial phenotypic switching [21, 22], stem cell differentiation [23], and immune cell lineage selection [24]. The first synthetic toggle switch was engineered in 2000 [8], and it’s often cited as a foundational paper in synthetic biology, which illustrates that the genetic circuits can be designed to control cellular behavior predictably. A toggle switch corresponds to *M* = 2 in Eq. (3) and *λ* = 0 in Eq. (2) with the Hill function [25]

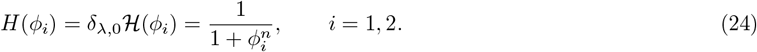

We also set the Hill coefficient *n* = 2. Equation (12) recovers the deterministic dynamics in the lowest order Ω^1*/*2^ as

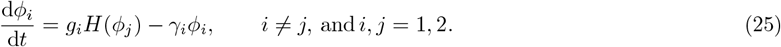

In this case, equating terms of Ω in the expansion, one obtains from Eqs. (16) and (17)

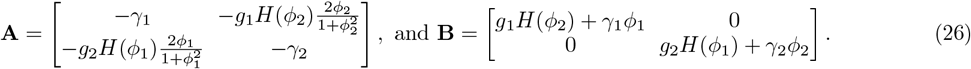

As shown in the general formalism, given by Eq. (18), being Gaussian, all higher moments of the variables, except for the first two, vanish. We are interested in the behavior of the non-vanishing moments only, namely, the mean and variance, which can be computed from Eqs. (20)(21). One finds for the first moments

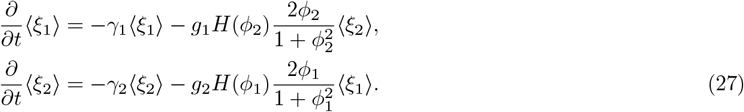

Similarly, for the second moment

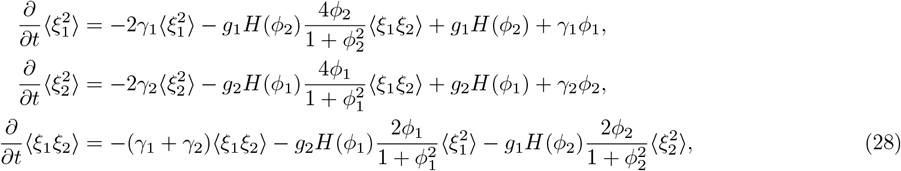

This gives us the auto- and cross-correlations in the intrinsic number fluctuation of the total number of type-A and type-B genes.

#### 1. Stationary-state fluctuations: First and second moments

Let us analyze the behavior of the mean fluctuations in the stationary state, ⟨*ξ*_1_⟩ ^*s*^ and ⟨*ξ*_2_⟩^*s*^. Note that in the stationary state, the variables *ϕ*_1_, *ϕ*_2_ take a stationary value, 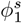 and 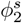, say, respectively. For Eq. (27) to have a non-trivial solution, one must have

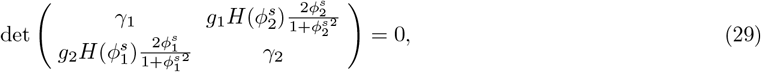

which simplifies

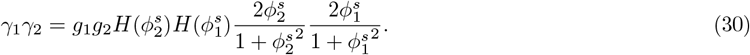

One has in the stationary state for the mean fluctuations

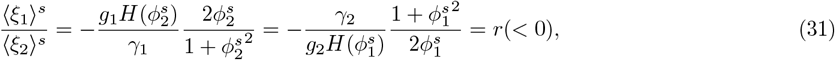

where *r* is a constant. This shows that the mean fluctuations are dictated by the interplay between the system parameters (growth and decay rates) as well as the stationary values of the variables. The negative sign implies an anticorrelation between them, meaning that an increase in one fluctuation corresponds to a decrease in the other. Otherwise, the trivial solution is

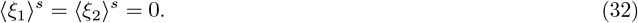

Now, for the variance, Eq. (28), in matrix form,

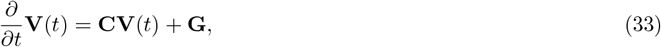

where

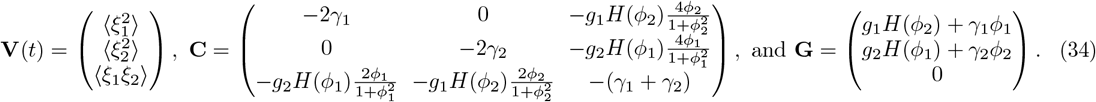

Since the coefficient matrix elements *C*_*ij*_ involve *ϕ*_1_ and *ϕ*_2_, which evolve in time, a full time-dependent solution to Eq. (33) is hard to obtain. Instead, we focus on the behavior in the stationary states attained at long times. Stationary-state variance of the fluctuations, which quantifies robustness of gene expression levels against intrinsic noise/perturbations, is thus given by

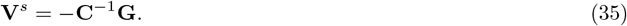

A measure of the strength of the fluctuation is given by the Fano factor, which for gene-A, as computed from Eq. (35), is given by

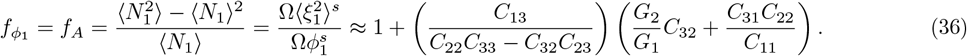

Similarly, the Fano factor for gene B is given by

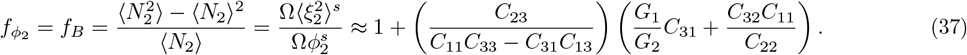

One can get an estimate of the order of magnitude of these quantities by substituting numerical values and calculating *C*_*ij*_*s*.

#### 2. Absence of inherent stochasticity in Hill functions

We now proceed to understand the crucial role played by inherent stochasticity in Hill functions in the stationary state. Had they been functions of average concentrations (macroscopic scenario) only (*ϕ*_1_ and *ϕ*_2_), the off-diagonal terms in **A** in Eq. (26) would vanish, yielding in the stationary state,

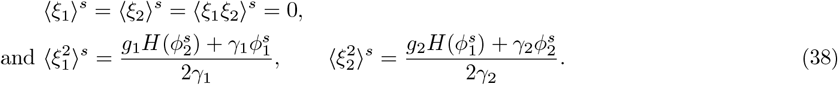

In such a case, one can further show that the FFs

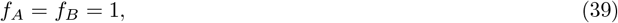

same as in the case of a Poisson distribution. The details are provided in the Appendix.

### B. Motif 2: Toggle switch with self activation (TSSA)

In this section, we study the impact of intrinsic noise in another motif, a variant of the Toggle switch. The motif involves two self-activation loops coupled with a toggle switch - Toggle switch with self-activation (TSSA). The deterministic stationary-state behavior of TSSA exhibits, in addition to the two expression levels of the toggle switch (high A-low B) and (low A-high B), a third stable steady state (intermediate A-intermediate B) in the system. The deterministic dynamics of a *M* -node GRN with self-activation follows the Eq. (1), with the only difference that the transition rate now follows

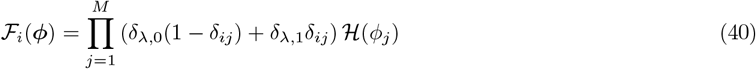

The mathematical framework discussed in Sec. III is also applicable in this case, and indeed,

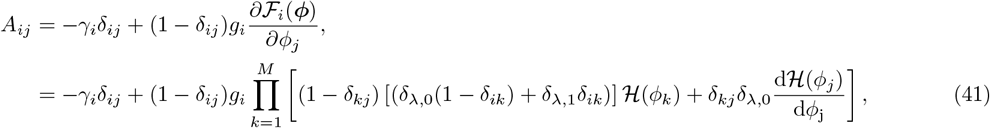

and the diagonal matrix ***B*** is given by

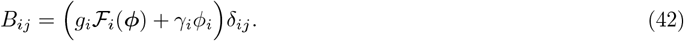

One thus obtains for TSSA with Hill coefficient *n* = 2 from Eqs. (41) and (42)

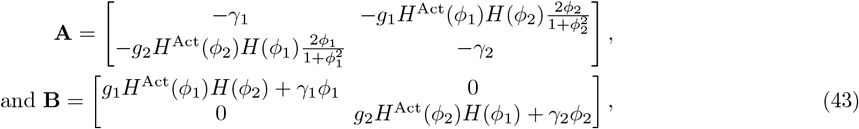

where *H*^Act^(*ϕ*_*i*_) = *δ*_*λ*,1_ℋ(*ϕ*_*i*_).

The mean and variance evolve in time, which can be computed using Eqs. (20)(21). For the first moments (mean)

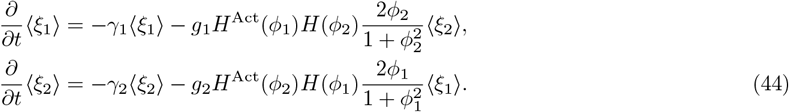

Similarly, for the second moment (variance)

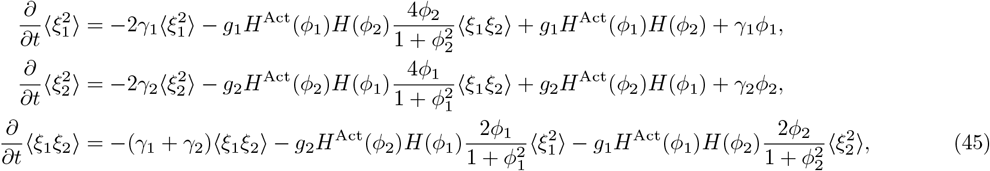

As in the Toggle switch case, here also, one can compute the stationary-state variance of the fluctuation, which is given by Eq. (35). Here, the matrices are given by

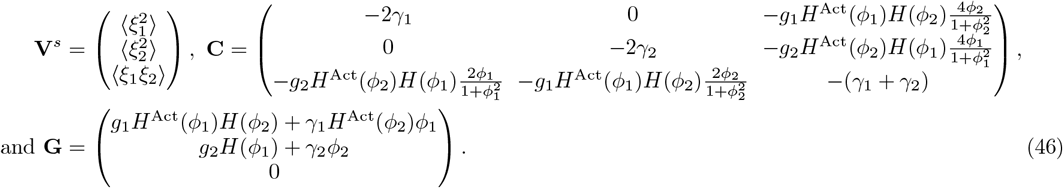

The FFs for genes A and B are given by Eqs. (36) and (37), with the matrix elements *C*_*ij*_ defined in Eq. (46).

## V. NUMERICAL RESULTS AND DISCUSSION

This section validates our theoretical approach to capturing the inherent stochasticity through the nonlinear coupling and low copy numbers of the genes at the microscopic level. To this end, we perform an *in silico* experiment of the system using two complementary approaches: the stochastic formulation of chemical kinetics by Gillespie (1977) [26], and, for the macroscopic limit, numerical integration of the corresponding deterministic dynamics (of ‘average’ concentration) using a fourth-order Runge–Kutta scheme with a time step *dt* = 0.001. As a representative case, we consider the simplest motif, the Toggle switch only. However, our theoretical results can easily be validated for other motifs with more complex emergent behaviors.

Here we present results on various possible steady states (fixed points) obtained by numerically integrating the deterministic dynamics, Eq. (25).

We first analyze the role of degradation rates *γ*_1_ and *γ*_2_ on the deterministic dynamics of TS, while keeping the production rates fixed at *g*_1_ = *g*_2_ = 1. To probe the stationary behavior, we initialize the system at a given state and let it relax to stationarity, in the limit *γ*_1_*/γ*_2_ → 0. We then adiabatically tune the ratio *γ*_1_*/γ*_2_ by increasing *γ*_1_ (with *γ*_2_ fixed) to large values and subsequently decreasing it along the same path. During this process, we monitor the stationary concentration of species *B*, denoted by *ϕ*_2_, as a function of *γ*_1_*/γ*_2_. The adiabatic variation ensures that the system remains close to stationarity throughout.

Figure 2 shows the variation of *ϕ*_2_ under forward and backward sweeps of *γ*_1_, for representative values of *γ*_2_ = 0.2, 0.3, 0.4, 0.5 and 0.6. For small *γ*_2_, the forward and backward trajectories differ, forming a hysteresis loop whose area gradually decreases as *γ*_2_ increases. Beyond a critical *γ*_2_, for instance, for *γ*_2_ = 0.6 the forward and backward curves coincide to numerical precision, indicating the disappearance of hysteresis. This behavior suggests a transition in the system’s response: for low *γ*_2_, the switch between low-*B* and high-*B* states is discontinuous, while at larger *γ*_2_ the transition becomes continuous.

**FIG. 2.**
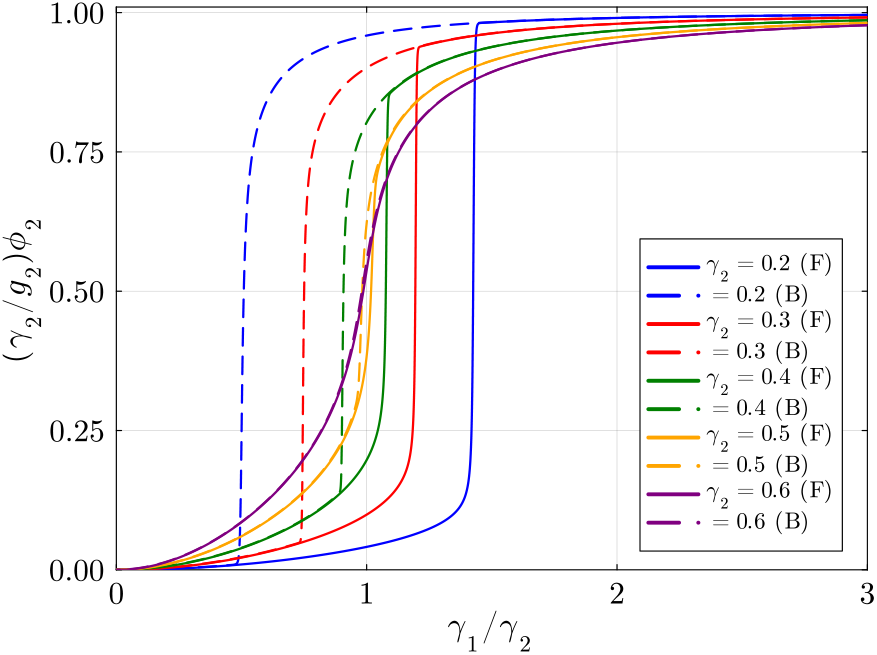
**TS**: Signature of monostability and bistability (hysteresis behavior) in the stationary-state dynamics of a TS obtained from its deterministic description, Eq. <u>(25),</u>as the parameter *γ*_1_ is tuned adiabatically keeping *γ*_2_ fixed at various values. Here, F and B denote forward and backward sweeps, respectively. The choice of parameters is: *g*_1_ = *g*_2_ = 1.0. (Similar complementary hysteresis behavior can be obtained for the A-type gene.)

**FIG. 3.**
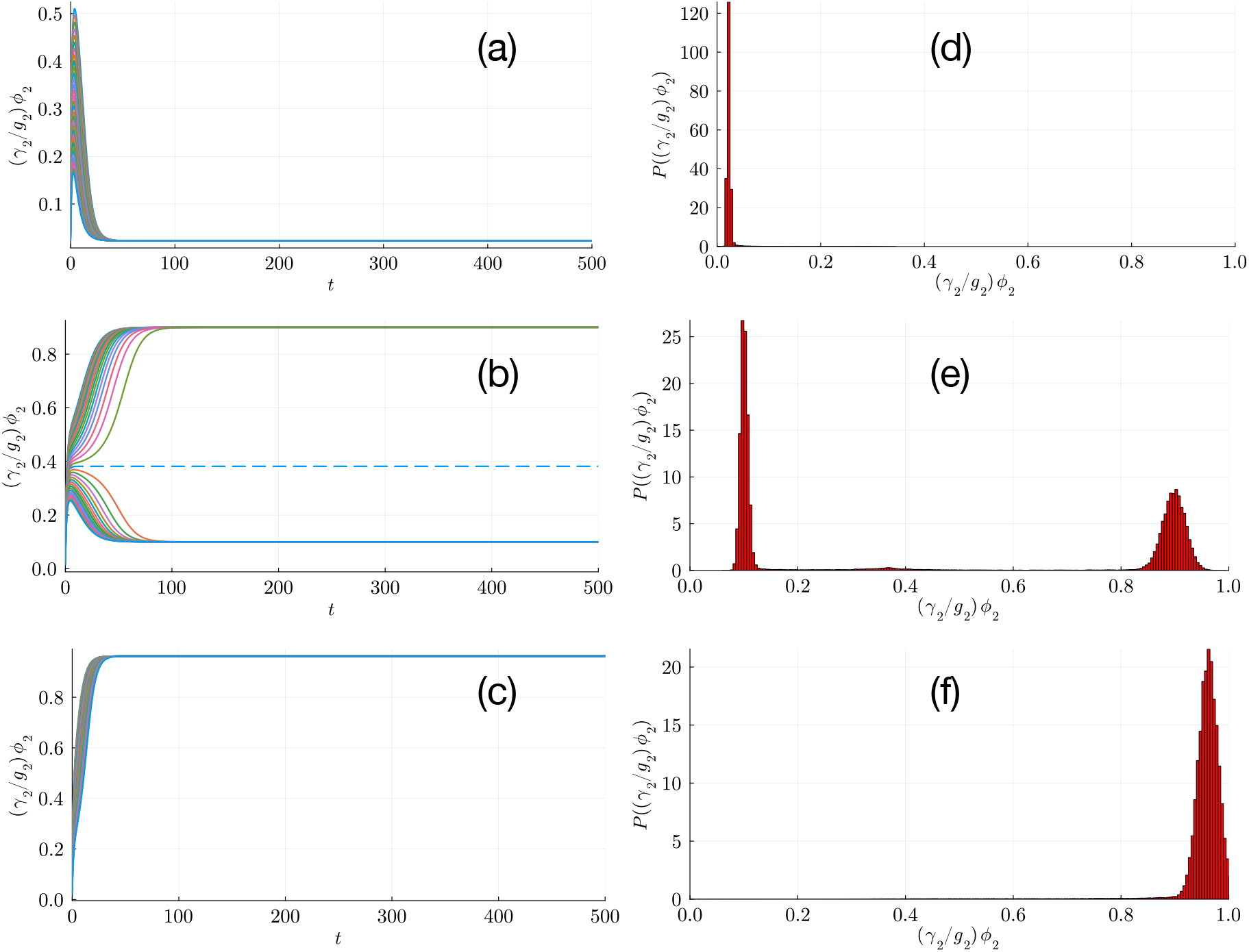
**TS**: Deterministic and stochastic dynamics of the Toggle switch at *γ*_2_ = 0.3, while *g*_1_ = *g*_2_ = 1.0. Panels (a–c) show *ϕ*_2_ trajectories for multiple initial conditions at *γ*_1_*/γ*_2_ = (a) 0.5, (b) 1.0, and (c) 1.5. Panels (d–f) show corresponding stationary distributions *P* (*γ*_2_*ϕ*_2_*/g*_2_) from Gillespie simulations. Monostable regimes [panels((a), (d)) and panels((c), (f))] display unimodal distributions at deterministic fixed points, while the bistable regime (b,e) exhibits a bimodal distribution, matching the two fixed points. The results highlight bistability and validate Eq. <u>(25),</u>the mesoscopic description obtained from LNA at its lowest order.

From a biological perspective, the presence of hysteresis corresponds to *cellular memory* : once the system switches to a high- or low-expression state, it resists switching back, thereby stabilizing a specific gene expression program [27]. Such bistability underlies critical decision-making processes in cell fate determination. The disappearance of hysteresis at larger *γ*_2_ implies a *loss of memory and increased tunability*, where gene expression levels can respond smoothly and reversibly to parameter changes. In the language of critical phenomena in statistical physics, this maps directly to phase transitions: the TS exhibits both *first-order transition* (corresponds to bistability with hysteresis) and continuous transitions between low and high expression states [28–30]. The disappearance of hysteresis at a critical *γ*_2_ can thus be interpreted as a *tricritical* -like point where these two regimes meet, in analogy with *tricritical* points observed in statistical physics systems [31–33].

With these fixed points in hand, we proceed to verify the mesoscopic description of the TS as obtained from the LNA. Fig. (3) consists of six panels: (a,b,c) show the time evolution of *ϕ*_2_ for three representative values of *γ*_1_*/γ*_2_, corresponding to three different dynamical regimes: *γ*_1_*/γ*_2_ = (a) 0.5, (b) 1.0, and (c) 1.5, with *γ*_2_ fixed at 0.3. Each panel displays trajectories from multiple initial conditions obtained by integrating the deterministic ODEs. Panels (a) and (c) demonstrate that all trajectories relax respectively to the low- and high-expression states, whereas panel (b) illustrates bistability: different initial conditions relax to distinct stable states. In all cases, after a transient period, trajectories converge to their corresponding deterministic fixed points.

The corresponding stochastic results are shown in panels (d,e,f), where we plot the stationary probability distribution *P* (*γ*_2_*ϕ*_2_*/g*_2_) obtained from Gillespie simulations for the same parameter values: *γ*_1_*/γ*_2_ = (d) 0.5, (e) 1.0, and (f) 1.5, with *γ*_2_ = 0.3. Panels (d) and (f) show unimodal distributions with peaks located at the deterministic fixed points [panels (a) and (c) respectively] within numerical accuracy of 10^−3^, consistent with the system being monostable in those regimes. By contrast, panel (e) exhibits a bimodal distribution with peaks centered very close to the two deterministic fixed points [panel (b)] within numerical accuracy of 10^−3^, reflecting the coexistence of two stable expression states. This demonstrates excellent agreement with Eq. (25), the lowest-order result of the LNA, which recovers the macroscopic deterministic dynamics, while also capturing the stochastic fluctuations around the fixed points.

We now quantify the amount of fluctuations in the stationary state of gene expression by computing the FF. Fig. 4 shows the FF for *A*-type [panels (a, c)] and *B*-type [panels (b,d)] genes as a function of *γ*_1_*/γ*_2_, for two fixed *γ*_2_ values corresponding to two distinct scenarios: a continuous transition for *γ*_2_ = 0.6 [panels (a,b)] and a discontinuous transition for *γ*_2_ = 0.3 [panels(c,d)], between the High-B (low-A) and Low-B (High A) states. In the continuous case, each type of gene exhibits monostability across the parameter range of interest, whereas in the discontinuous case, bistability emerges only in certain regimes, with monostability elsewhere, see Fig. 2.

**FIG. 4.**
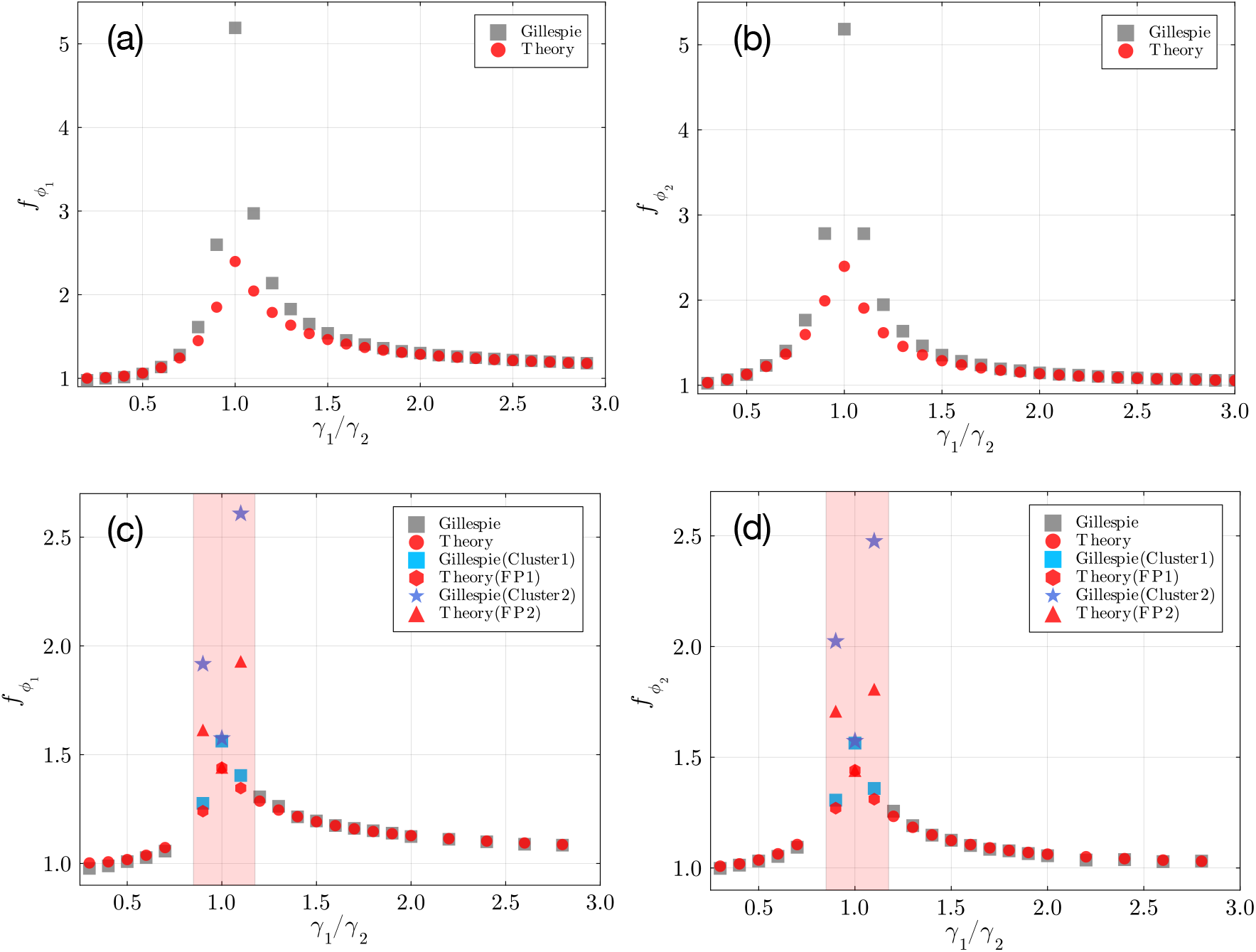
**TS**: Fano factors (FF) for *A*-type [panels(a, c)] and *B*-type [panels (b, d)] genes as a function of *γ*_1_*/γ*_2_. Panels (a, b) correspond to *γ*_2_ = 0.6 (continuous transition), and panels (c, d) correspond to *γ*_2_ = 0.3 (discontinuous transition). Gray-filled squares denote results from Gillespie simulations, averaged over multiple trajectories and initial conditions, while red-filled circles represent theoretical predictions from the LNA in the monostable region. The shaded region indicates the region of bistability. In the bistable region, distributions (of each gene type) show two peaks (clusters), with theoretical FFs corresponding to each deterministic fixed point labeled as FP1 and FP2. Good agreement between simulations and theory is observed in the monostable region (away from the transition point) and deep in the bistable regions, whereas deviations occur near the bifurcation points due to enhanced fluctuations. Other parameters: *g*_1_ = *g*_2_ = 1.0.

In the monostable regime, multiple stochastic trajectories are generated using Gillespie simulations starting from different initial conditions. Two types of averaging are performed: (i) over many realizations for a given initial condition, and (ii) over different initial conditions. For each initial condition, we compute the variance-to-mean ratio from the corresponding ensemble of trajectories, which yields a distribution across many initial conditions. The Fano factor is then computed as the mean of the distribution, and is shown as gray-filled squares in Fig. 4.

In the bistable regime (shaded region), the distributions of type-*A* or type-*B* genes exhibit two distinct peaks, obtained from the ensemble of the stochastic trajectories generated over many realizations and many initial conditions, which we label *cluster 1* and *cluster 2*. The variance around each peak is computed separately, and the corresponding variance-to-mean ratios yield the FFs for cluster 1 and cluster 2, respectively.

These results are compared with theoretical predictions obtained from the LNA. For each parameter set in the monostable region, the deterministic fixed points are computed from the ODE dynamics, and the corresponding FFs 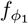 and 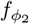 are calculated using Eqs. (36) and (37) for *A*-type and *B*-type genes, respectively. These theoretical predictions are shown as red filled circles. In the bistable case, the theoretical FFs are estimated using the two corresponding deterministic fixed points. In Fig. (4), Theory (FP1) and (FP2) indicate the theoretical FFs for cluster 1 and cluster 2, respectively.

The comparison shows good agreement between simulations and theory in the monostable regime and deep in the bistable regime, with deviations near (i) the transition region between the low and high states in the continuous case, and (ii) the boundary between the monostability and bistability. The fluctuations are enhanced in these regions, and the LNA becomes less accurate. Note that, for small values of *γ*_1_*/γ*_2_, the numerically computed FFs 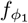 and 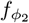sometimes fall below unity. In principle, the minimum value should be unity (Poissonian case). This is likely due to low copy numbers, small basal expression levels, and finite simulation times, which can underestimate the variance in the stochastic trajectories.

The fact that the LNA underperforms in the aforementioned cases is intuitive, especially from the perspective of the theory of phase transitions and critical phenomena in statistical physics. For continuous transitions, fluctuations of the order parameter grow strongly in the vicinity of the critical point and diverge exactly at criticality, which is beyond the applicability of the LNA [30, 34, 35]. Here, the fluctuation data from simulations, which show a crossover rather than a divergence, are attributed to the typical finite-size (here, finite cell volume) rounding off of a continuous transition [30]. For the discontinuous transition, this deviation can be understood in terms of finite-size effects. A discontinuous transition becomes sharp only in the thermodynamic limit (limit of infinite system size), where fluctuations are negligible except exactly at the transition point [30, 36]. In our present context, in a cell of finite volume, limited copy numbers of genes amplify intrinsic fluctuations near bifurcations, leading to departures from LNA predictions and occasionally triggering stochastic switching even before the deterministic threshold is reached.

### B. Variation with *g*_1_, *g*_2_

We next analyze the effect of varying the production rate ratio *g*_1_*/g*_2_ on the dynamics of the TS, keeping *γ*_1_ = *γ*_2_ = 1.

Fig. 5 shows the stationary concentration of species *B, ϕ*_2_, as *g*_1_ is tuned adibatically for fixed *g*_2_ = 1.0, 2.0, 3.0, 4.0, 5.0. For larger values of *g*_2_ (2≤ *g*_2_ ≤5), forward and backward sweeps of *g*_1_ exhibit hysteresis loops, indicative of bistability. The width of the hysteresis loop decreases as *g*_2_ is reduced. At *g*_2_ = 1.0, the forward and backward trajectories coincide, signaling a continuous transition between high and low-expression states. This behavior is similar to what we have previously observed in the case of dependence on degradation rates, with hysteresis corresponding to cellular memory and its disappearance indicating increased tunability.

**FIG. 5.**
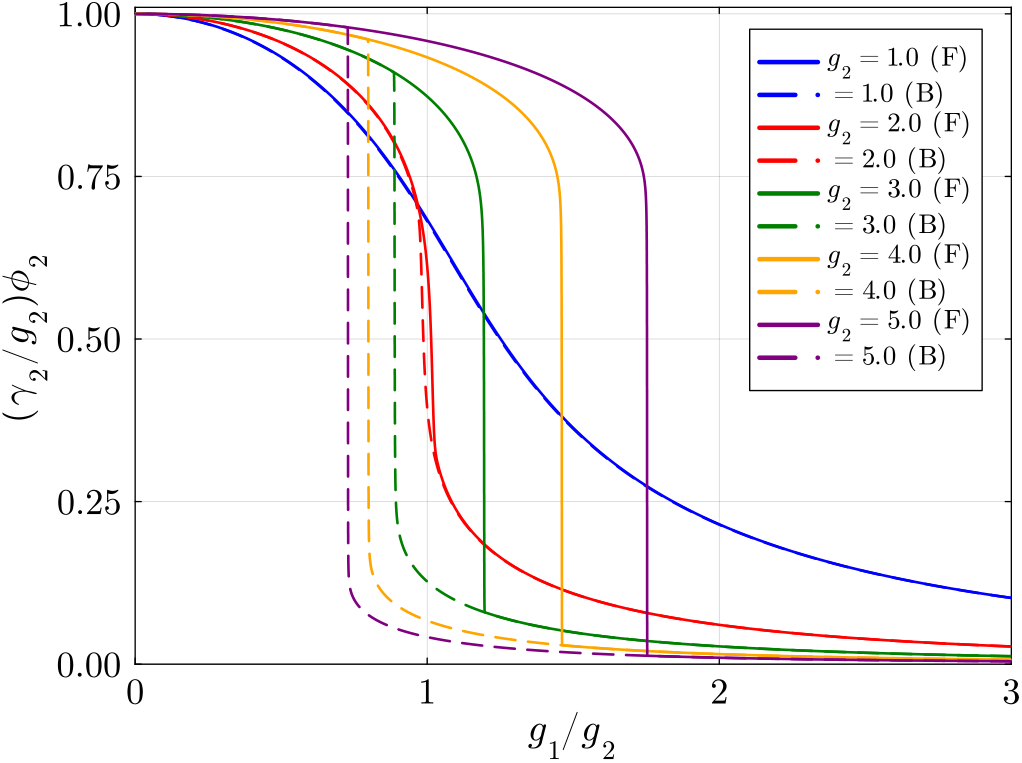
**TS**: Signature of monostability and bistability (hysteresis behavior) in the stationary-state dynamics of a TS obtained from its deterministic description, Eq. <u>(25),</u>as the parameter *g*_1_ is tuned adiabatically keeping *g*_2_ fixed at various values. Here, F andB denote forward and backward sweeps, respectively. The choice of parameters is: *γ*_1_ = *γ*_2_ = 1.0.

**FIG. 6.**
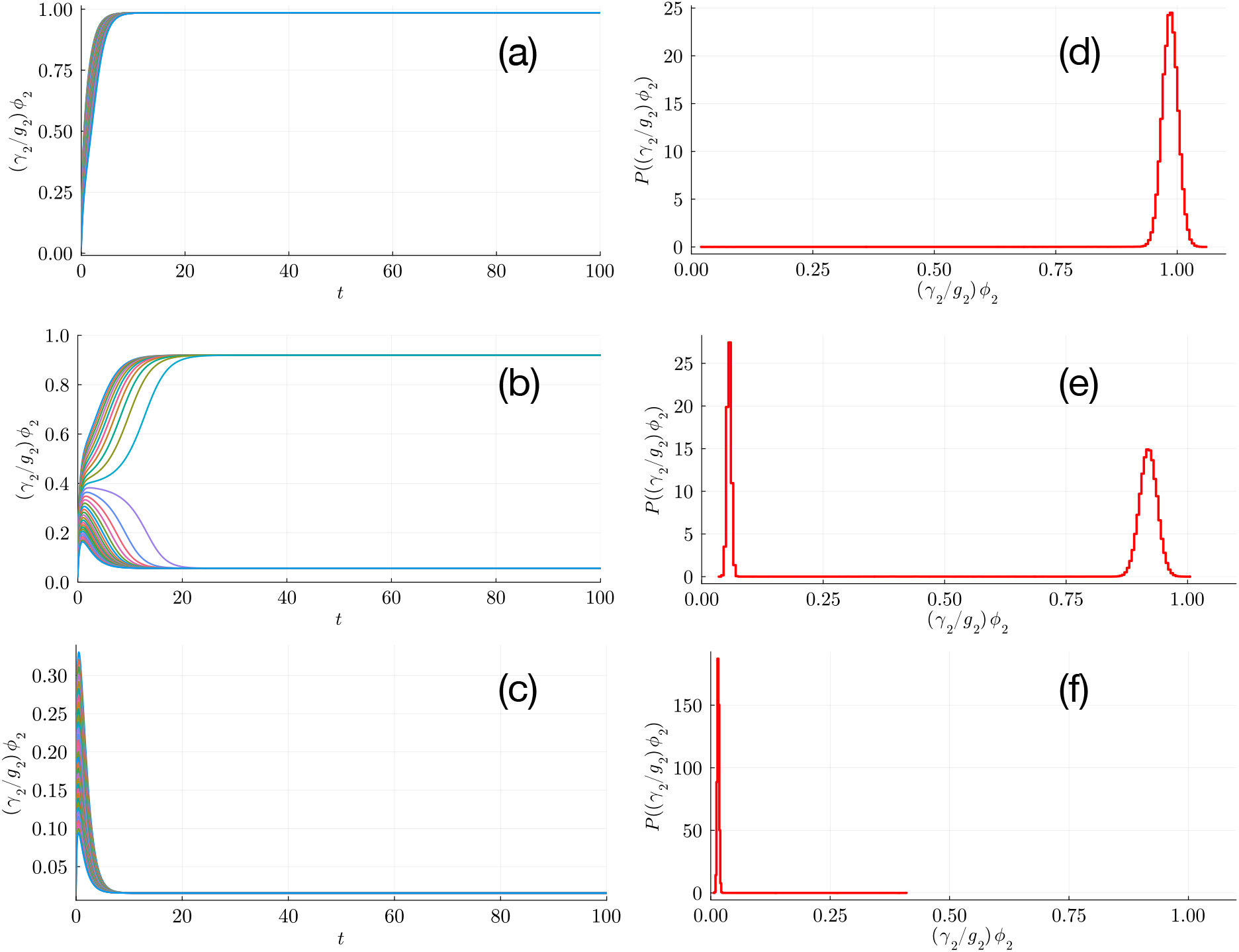
**TS**: Deterministic and stochastic dynamics at *γ*_1_ = *γ*_2_ = 1.0, while varying *g*_1_*/g*_2_. Panels (a–c) show *ϕ*_2_ trajectories for multiple initial conditions at *g*_1_*/g*_2_ = 0.5 (a), 1.0 (b), and 2.0 (c). Panels (d–f) show the corresponding stationary distributions *P* (*γ*_2_*ϕ*_2_*/g*_2_) from Gillespie simulations. Monostable regimes [panels (a,d) and panels (c,f)] display unimodal distributions at deterministic fixed points, whereas the bistable regime (b,e) exhibits a bimodal distribution, matching the two fixed points. These results highlight bistability and validate Eq. <u>(25),</u>the mesoscopic description obtained from the LNA at its lowest order.

**FIG. 7.**
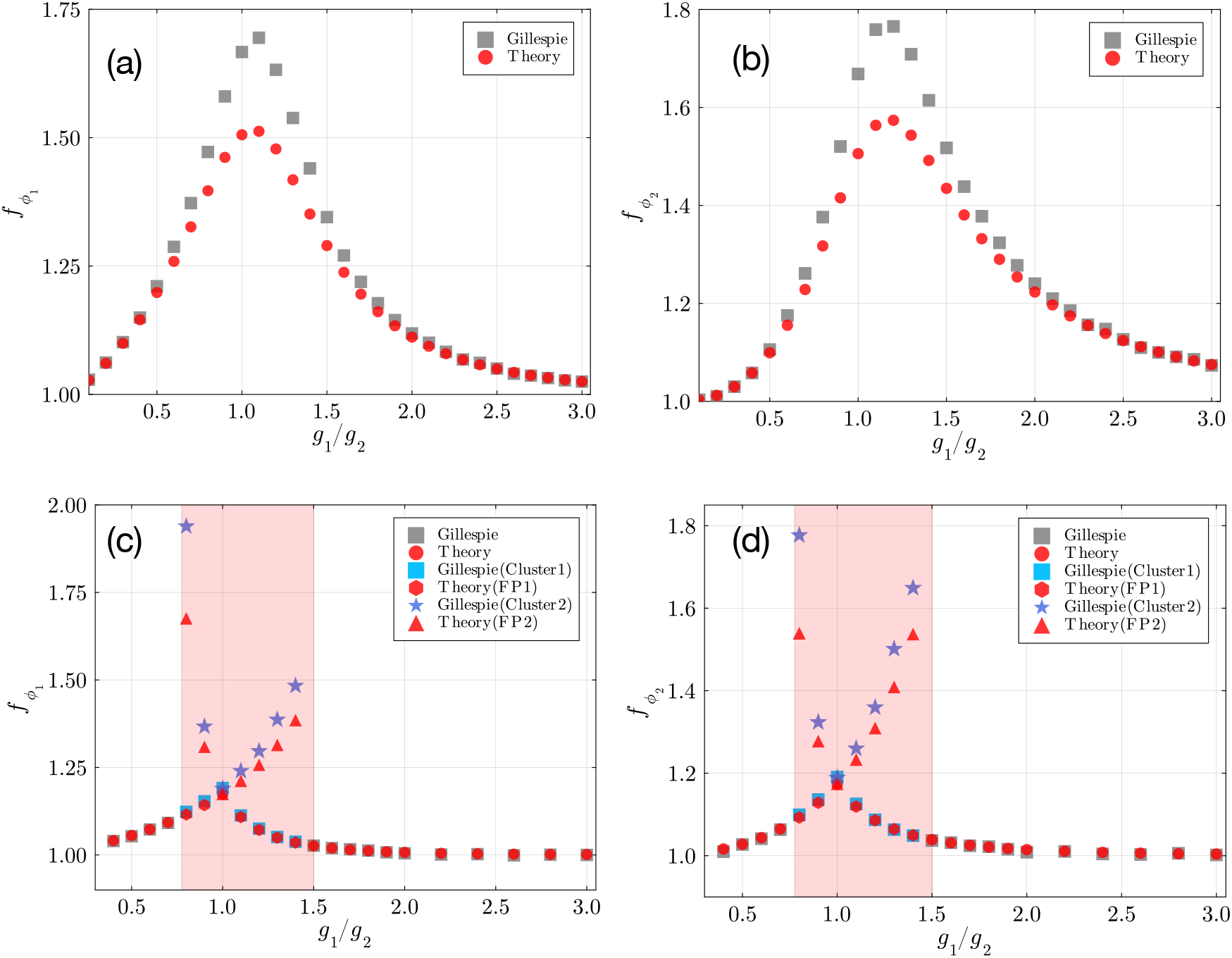
**TS**: The Fano Factors (FF) for *A*-type [panels (a,c)] and *B*-type [panels (b,d)] of genes, as a function of *g*_1_*/g*_2_. Panels (a, b) correspond to *g*_2_ = 1.0 (continuous transition), and panels (c, d) correspond to *g*_2_ = 5.0 (discontinuous transition). Gray-filled squares denote results from Gillespie simulations, averaged over multiple trajectories and initial conditions, while red-filled circles represent theoretical predictions from the LNA in the monostable region. The shaded region indicates the region of bistability. In the bistable region, distributions of the genes show two peaks (clusters), with theoretical FFs corresponding to each deterministic fixed point labeled as FP1 and FP2. Good agreement between simulations and theory is observed in the monostable region (away from the critical point of continuous transition) and deep in the bistable regions, whereas deviations occur near criticality (bifurcation points) due to enhanced fluctuations. Other parameters: *γ*_1_ = *γ*_2_ = 1.0.

Time evolution and stationary distributions from Gillespie simulations confirm these regimes, see Fig. (6). For representative values *g*_1_*/g*_2_ = 0.5, 1.0, 2.0 with *g*_2_ = 4.0, the deterministic ODE trajectories [panels (a–c)] show relaxation to either high or low-expression states, or bistability for intermediate ratios. The stochastic stationary distributions [panels (d–f)] reproduce unimodal or bimodal profiles consistent with the deterministic fixed points.

FFs, as shown in Fig. (7), depict two scenarios. For *g*_2_ = 1.0, the system undergoes a continuous transition [panels(a, b)] and remains monostable, while for *g*_2_ = 5.0 [panel(c,d)], a discontinuous transition leads to bistability in certain regimes. In monostable regions, as before, FFs are computed by averaging variance-to-mean ratios over multiple stochastic trajectories and initial conditions, whereas in bistable regions, FFs are obtained separately for each peak (*Cluster1* and *Cluster 2*). LNA predictions show agreement with that obtained from simulations, except near the transition point between the two gene-expression states (continuous case) and near monostable-bistable boundaries (in the discontinuous case). Qualitatively similar (as observed in the continuous transitions in the monostable regime of TS as depicted in Figures 4 (a, b) and (7) (a, b)) divergence of intrinsic noise (variance) has been observed in a fundamentally different biological context: conformational receptor dynamics near the ultrasensitive critical transition point, when an intracellular enzyme concentration in the *E. coli* signaling cascade is tuned [37, 38]. The system exhibits zero-order ultrasensitivity, in contrast to the multistable dynamics exhibited by GRN motifs discussed in the current study. While for the continuous case, the enhanced fluctuations near criticality (critical point of transition) reduce LNA accuracy, deviations near discontinuous transitions arise from intrinsic fluctuations in finite systems near the bifurcation points [30, 34–36].

Thus, the TS exhibits monostable and bistable behavior as the production and degradation rates are varied. The fluctuations around them are well-captured by LNA except near the monostable-bistable boundary (discontinuous case), and near the critical point of transition (continuous case), where enhanced fluctuations are expected from the theory of critical phenomena.

These findings thereby highlight the universality of phase transitions across diverse biological systems, and the examples vary from multistable gene regulatory dynamics and cell fate determination [39, 40] to metabolic coordination [41], motility of cells in a tissue [42], population dynamics, evolutionary transitions [43, 44] demonstrating that similar critical phenomena emerge across vastly different biological scales and mechanisms [45, 46].

## VI. SUMMARY AND CONCLUSION

How can we understand nonlinearity-driven stochasticity in GRN motifs? Our present study develops a comprehensive theoretical framework to systematically investigate the role of inherent stochasticity in a nonlinearly coupled *M* -gene network motif. We observe that the LNA captures well the propagation of noise through the nonlinear cooperative interactions embedded in the Hill-type regulatory functions. This has been validated by comparing the lowest-order LNA, which recovers the deterministic dynamics, and the next-order LNA expressions, which provide the mean and variance of fluctuations around macroscopic fixed points, with stochastic simulations using the Gillespie algorithm as well as with solutions of the deterministic ODEs. We have obtained an estimate of the magnitude of fluctuations around each deterministic fixed point directly from the values of the fixed points obtained from the ODE solutions.

We explicitly analyzed the toggle switch motif and established a parallel with concepts from statistical physics, showing that it exhibits both first-order and continuous transitions, with gene concentration as an order parameter, between low- and high-expression states, with the existence of a tricritical-like point. We have demonstrated that the LNA provides good agreement with stochastic results in the monostable regime and deep within the bistable regime, provided the system is far away from the criticality (bifurcation point) for both types of transitions. Near criticality, i.e., in the vicinity of the bifurcation point, the LNA underperforms. We rationalize this deviation by borrowing concepts from critical phenomena in statistical physics, where fluctuations become strongly enhanced near critical points of continuous transitions and finite-size effects are amplified near first-order transition points, while at criticality, fluctuations diverge.

Biologically, this highlights that caution is needed when operating synthetic gene circuits near critical regimes, where stochastic effects can be strongly amplified, far exceeding the predictions of the LNA, potentially disrupting the intended network function [47]. Importantly, our approach provides a practical tool for synthetic gene regulatory network design, as the magnitude of noise can be estimated in advance using the deterministic fixed point values from ODE solutions, allowing designers to anticipate fluctuations and optimize circuit robustness before implementation.

While our analysis demonstrates the usefulness of the LNA in capturing intrinsic stochasticity in gene regulatory motifs, a more complete framework is needed to describe the enhanced fluctuations that arise near critical regimes. We have not included higher-order corrections in the system-size expansion. Previous works [47, 48], for instance, through effective mesoscopic rate equations [47] and tools such as iNA [48], have demonstrated that such corrections can be crucial for capturing stochastic dynamics in regimes where LNA underestimates fluctuations, particularly in oscillatory systems. Incorporating these higher-order terms, or comparing our results against such methods, remains an important next step.

Furthermore, our study has primarily focused on intrinsic noise under symmetric parameter choices. In real cellular systems, however, extrinsic noise and parameter heterogeneity, arising from cell-to-cell variability in gene expression rates, degradation rates, and other biochemical parameters, play a very crucial role that may alter the results [15, 49, 50]. Finally, we have concentrated on the toggle switch motif as a representative case. Extending our framework to larger and more complex circuits, such as genetic oscillators or feed-forward loops, would provide a more comprehensive validation.

While we have discussed the equivalence of the LNA to an Ornstein–Uhlenbeck (OU) description, we did not explicitly compare our results with Langevin simulations incorporating Gaussian white noise. Performing such comparisons will be an important step in fully benchmarking the framework. It is also worth mentioning that, although our predictions can in principle be directly compared with experimental measurements of noise in synthetic circuits, such a comparison was not carried out in the present study. Establishing this link to data will be essential for quantitatively benchmarking the accuracy of LNA-based predictions and for demonstrating their practical value in circuit design [51]. Nevertheless, our framework already offers predictive power for synthetic biology. Estimating the magnitude of noise directly from deterministic ODE solutions enables the identification of parameter regimes where synthetic circuits are likely to remain robust or fragile to stochastic fluctuations. To this extent, our results provide a rigorous mathematical foundation for developing design principles that aim to construct reliable and tunable GRN motifs capable of self-organizing into multiple, distinct, coexisting stable states and persistent rhythmic patterns.

## DATA AVAILABILITY

The data that support the findings of this study are available from the first author upon reasonable request.

## ACKNOWLEDGMENT

M.S. acknowledges support by the Deutsche Forschungsgemeinschaft (DFG, German Research Foundation) under Germany’s Excellence Strategy EXC 2181/1-390900948 (the Heidelberg STRUCTURES Excellence Cluster). U.R. acknowledges the support of DST-INSPIRE, India (Sanction Letter No. DST/INSPIRE/04/2022/003052 dated 27.02.2023).

## AUTHOR CONTRIBUTIONS

M.S. and U.R. developed the idea and designed the problem. M.S. executed the analytical and numerical analyses. Both M.S. and U.R. discussed the results and wrote the manuscript. The work was carried out independently by both authors.

## Appendix A: Stationary-state fluctuations for a Toggle switch

In this section, we provide explicit calculations of the FFs derived from the stationary-state fluctuations for a toggle switch (TS). As provided in Eq. (35) in the main text, the stationary-state variance of the fluctuations for a TS, in matrix form, is given by

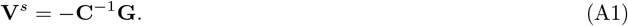

Explicitly,

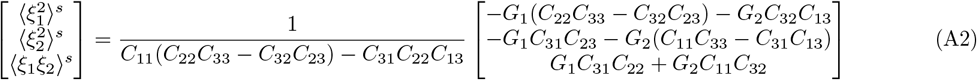

where the elements *G*_*i*_ and *C*_*ij*_ are given by Eq. (34). The Fano factor for gene-A is given by

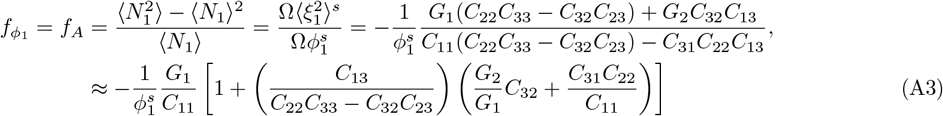

One immediately identifies 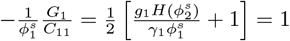. Thus, we obtain the Fano factor for gene-A as

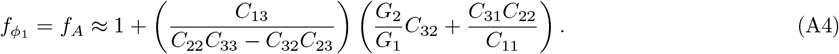

Similarly, the Fano factor for gene B is

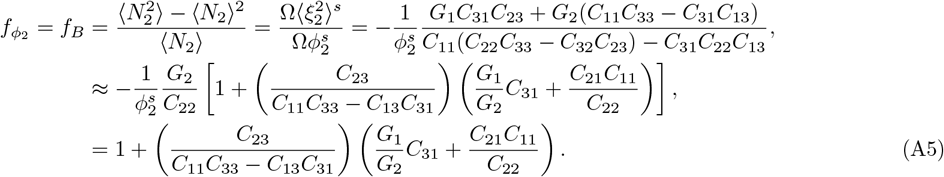

